# Social Approach Dynamics in Freely-Moving Rat Dyads are Not Impacted by Lesions to the Retrosplenial Cortex

**DOI:** 10.64898/2026.10.19.760292

**Authors:** Mckenna I. Lowrie, Jaidyn E.M. Smith, Angela T.N. Le, Diano F. Marrone, Nathan Insel

## Abstract

Social interaction in rodents can involve a dynamic interplay of approach and withdrawal movements, requiring the brain to rapidly update information about where others are and where they may intend to go. In mammals, the posterior parietal cortex (PPC) and adjacent, reciprocally-connected retrosplenial cortex (RSC) are high-level hubs for spatial information processing. To test the role of these regions in dynamic social approach, we evaluated rats’ relative orientation, movement, and position (ROMP) with a freely-moving social-stimulus rat following bilateral lesions to the PPC or anterior RSC. RSC animals showed overall more and faster movements, and greater attention to environment boundaries across multiple behavioral conditions (open field, social, object, static social, and dynamic object) compared with sham-lesioned controls. Increased attention to boundaries was correlated with reduced orienting to a social stimulus; however, this appeared to have little observable impact on the social interactions themselves. Rear-sniffing was lower in RSC rats, but only during the latter half of the 10 minute social exposures, and there were no detected differences in the dynamics of their social approach compared with sham controls. PPC lesion spread was more limited, and social behavior was mostly indistinguishable from sham controls, though this group was more likely to chase the dynamic object. Results suggest that social approach within short (peripersonal) distances is largely accomplished through sensorimotor adjustments that do not require higher-level, spatial-predictive processing taking place in the RSC.

## INTRODUCTION

An animal’s ability to track the position of others is essential for predator-prey and social interactions, but also carries heavy computational demands. In contrast with self-movement, the movements of others can be unpredictable and require observers to rapidly update inferences about space, and make inferences about others’ intentions. Spatial information is also complex, as it can include where others are relative to an observer as well as where they are relative to environmental stimuli, along with often important details about how others are oriented or moving through space. How all of this information is integrated in the rodent brain is largely unknown—even to the point that we lack a clear idea of where, anatomically, to look.

Research in human and non-human primates have offered valuable insight into the functional neuroanatomy of spatial as well as social information processing. Early work— which largely took a perception-to-cognition (“outside-in”) approach, revealed the parietal cortex as critical for spatial cognition and behavior (Mishkin, 1972; Mishkin et al., 1983). Dorsal areas of the temporal lobe, including the superior temporal sulcus and adjacent temporal-parietal junction, have recently been shown to use visual movement information to process social intentions (reviewed by Pitcher & Ungerleider, 2021). The parietal cortex itself may include social information in the form of agent-independent action representations (i.e., “mirror neurons”, sensitive to both self- and other-initiated actions; Fogassi et al., 2005). All of these pathways have been examined relatively extensively in human neuroimaging, but have received less attention in rodent research, leaving gaps in our understanding of how they map onto the cortical architecture across mammalian taxa.

In contrast to primates, research on spatial processing in the rodent brain has historically followed a more “inside-out” path. Early studies revealed the dorsal hippocampus to contain map-like patterns of navigational space (O’Keefe & Dostrovsky, 1971; O’keefe & Nadel, 1978). Subsequent work demonstrated a role of the medial entorhinal cortex for the stability and partial generation of this space (Fyhn et al., 2004; Hafting et al., 2005; Hales et al., 2014; Igarashi, 2016; Jacob et al., 2020; Miao et al., 2015), and a role of the retrosplenial cortex (RSC) in path integration behaviors, potentially by bridging “self-“ (egocentric) and “environment-centered” (allocentric) reference frames (Alexander et al., 2023; Alexander & Nitz, 2015; Cooper et al., 2001; Mao et al., 2020). The role of posterior parietal cortex (PPC) in navigation has also been examined, which is interconnected with the RSC and is a site where information about goals, movement, and egocentric space converge (Alexander et al., 2022; Nitz, 2006; Whitlock, 2017; Whitlock et al., 2012; Wilber et al., 2014). The PPC also contributes to hippocampal representations of space (Save et al., 2005).

Few studies have evaluated the role that these spatial processing regions play for social-spatial behavior and cognition. Recordings from the hippocampus in both rats and bats have revealed the hippocampus can show “mirror-like” place selectivity when animals observe others moving through the environment (Danjo et al., 2018; Omer et al., 2018), and signal information about social-context (Forli & Yartsev, 2023; Ray et al., 2025). There are also reasons to believe the PPC may take part in social-spatial updating. In one study, neurons from the PPC were recorded as rats chased the light from a laser pointer held by an experimenter. Although not explicitly “social” (in that the experimenter’s laser movements may not have been perceived by rats as a social stimulus) this offered evidence that neurons in the region were often responsive to a dynamically-moving goal stimulus. Whether the PPC is *necessary* for a mammal to make precise movements to a dynamically moving goal has not been as well studied—though the idea is supported by indirect evidence from studies of reaching behavior in humans with damage to the parietal cortex (Gréa, 2002; Pause et al., 1989; Pisella et al., 2000). It is also possible that whole-body movements in relation to a social stimulus rely on the RSC, which in rodents is known to be extensively, reciprocally connected with the PPC (Glickfeld & Olsen, 2017; Mesina et al., 2016; Oh et al., 2014; Olsen & Witter, 2016; Wang et al., 2012; Wilber et al., 2015; Zingg et al., 2014). The rodent RSC is relatively large with many subdivisions, and recent evidence points to the region contributing to some autism-like behaviors—including reduced investigation of social novelty—found in certain genetic mouse models (Garrido et al., 2022; Shang et al., 2021; Yang et al., 2021). Furthermore, it has been suggested that the RSC could support functions linked to medial-wall parietal regions in humans (Xiang et al., 2023). One medial parietal region, the precuneus, has been shown to increase activation during social observation, and may track social expectations (Petrini et al., 2014). Here we test the hypothesis that lesions to either the PPC or the RSC alter how rats approach one-another and engage in social interactions. The RSC and PPC may complement one-another with differential contributions to processing sensory-based egocentric information (perhaps higher in PPC) relative to allocentric information (perhaps higher in RSC). But are together part of an integrated system for spatial cognition (Alexander et al., 2023; Clark et al., 2018), and may be critical for updating movement decisions in an environment with dynamic, social stimuli.

A challenge for studying the role of brain regions in social approach is that there are no consensus measures for the complex dynamics of social movement. A key principle underlying social interaction is that it is uncertain: socially-directed movements may be altered and updated according to uncertain changes in another animal’s pose, trajectory, or vocal signaling. One-dimensional behavioral tests restrict the degree of uncertainty, limiting assessment of how relative distance and movement are regulated and, correspondingly, evaluations of impacts of brain manipulations on social behavior. In the present work we specifically examine how patterns of behavior, including socially-directed movements (those immediately prior to an identified interaction), were impacted by lesions to the PPC and RSC. To do this, we derive measures of relative orientation, movement, and position (ROMP) from multi-animal pose-estimation obtained using recently-developed computer vision methods (Lauer et al., 2022; Robinson et al., 2025). Based on the hypothesis that both the PPC and RSC process social space, we predicted that approach would more often include overly rapid (high velocity and acceleration/deceleration) and intrusive (approach ending in close touch), or, alternatively, overly slow/cautious (slower acceleration/deceleration, potentially ending with animals still farther apart) movement with a peer or moving object.

## METHOD

### Subjects

Thirty-seven female Sprague Dawley rats (Charles River, Quebec) were used as experimental animals and an additional 8 female rats were used as social stimulus animals for this study. Numbers for each lesion group were determined based on experimental power of interaction level differences observed in past, behavioral manipulation studies. No animals were excluded, though a prior experiment helped establish the lesion protocol. Experimental rats weighed 250-350g and were four to six months old when behavioural training began; stimulus animals were between 3 to 5 months old. Only females were used to reduce the possibility of fighting during social exposures. Rats were housed in pairs on a 12-hour reverse light cycle. All procedures were in accordance with and approved by the Wilfrid Laurier University Animal Care Committee.

### Surgery

Each lesion rat was anesthetized with isoflurane and received bilateral infusions of 10 mg/ml ibotenic acid in phosphate buffered saline. One group of rats received 5 infusions in each hemisphere to the PPC (PPC group; n = 11), another received 6 infusions in each hemisphere to the RSC (RSC group; n = 11). Infusions were performed by a microinfusion pump (KD Scientific Holliston, MA) at a constant rate of 0.1 µL/min, up to a volume of 0.10 µL. In control animals, rats received the same anesthesia and scalp incision without craniotomy (sham group; n = 15). The reason for the higher number of sham animals is that the study was performed in 4 rounds of surgeries; one round included only PPC rats (each with a paired sham), and one round included only RSC rats (also with paired shams). Statistical comparisons evaluated each lesion group separately against the 11 controls they were paired with. Following surgery, rats were single-housed and given at least 6 days to recover before experimental testing began.

### Apparatus & Testing

All behavioral tests took place in a 50 x 50 cm chamber with 50 cm high walls and recorded at 60 frame/s from an overhead Logitech Brio 4K Pro webcam (Logitech; Lausanne, Switzerland). Rats were tested one time each day, with each test using a different exposure profile that always followed the same sequence: **(1) Open field**. To test general measures of movement and spatial exploration, the rat was placed in the empty, unfamiliar chamber alone for 10 min. **(2) Social**. To evaluate the effects of lesions on social approach, the rat was place in the chamber with a same sex, unfamiliar stimulus rat and allowed to interact freely for 10 min. The rear of one of the two animals was marked with an animal marker (Stoelting Co; Wood Dale, IL) to distinguish them during data processing. **(3) Static social**. To control for the effects of social stimuli that are unrelated to dynamic movement, the rat was placed for 10 min into a chamber where a new, unfamiliar peer stimulus rat was confined within a polycarbonate cylinder (15.5 cm diameter x 45 cm height). One side of the cylinder had slits cut to allow animals to sniff one-another. **(4) Object**. The rat was placed in a chamber where a single object sat at the chamber center for 5 min. Here, 5 minutes was used instead of 10 due to prior, pilot tests showing an absence of movement activity after 5 min. Since primary analyses considered movement around investigative actions separately across epochs, rather than comparing interaction levels between epochs, reducing time to omit inactivity did not introduce bias in any statistical comparison. **(5) Dynamic object**. As another point of comparison for dynamic stimulus movement, without other social cues, rats were placed for 5 min in a chamber with a small, remote controlled 40 mm diameter “Sphero Mini” ball (Sphero Inc.; Boulder, CO), which was then moved around remotely by an experimenter. The experimenter was told to move the robot around the chamber, but also to approach the rat slowly to encourage investigation and chasing. All conditions are illustrated in **Figure 1**.

**Figure 1.**
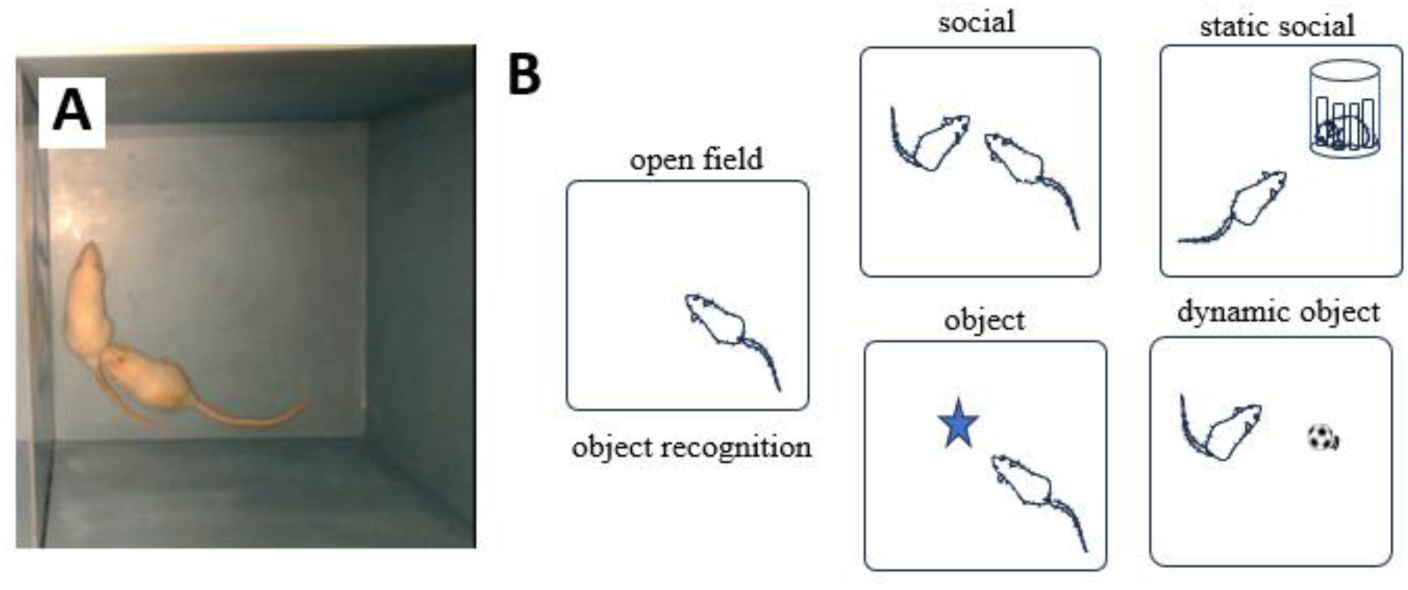
Apparatus and conditions. *Note*. (A) A screenshot depicting two rats interacting in the experimental chamber. (B) The exposure conditions, sequenced from first to last (top row left to right to bottom row left to right). Description of each condition in Methods.

### Histology & quantification of lesion

Within five days of the final behavioural recording, rats were euthanized with an intraperitoneal injection of a 1:1 ratio of sodium pentobarbital, (2.5 mL; 240 mg/ml) and Xylocaine (2.5mL; 20mg/mL) and perfused intracardially with formalin for tissue fixation. Tissue was cryoprotected by storing in 30% sucrose for one or more days, then sectioned at 50 μm. To assess lesion placement and extent, sections were stained with cresyl violet and verified under a microscope. Volumetric comparisons to healthy tissue from a rat brain atlas (Paxinos & Watson, 2007) was used to estimate the percent of damage to the target and off-target regions. SketchAndCalc Area Calculator (Palm Coast, FL) was used to calculate the area of the lesion as well as the area of remaining intact tissue in the target region.

### Position Tracking

Pose estimation was completed for each video with DIPLOMAT (Robinson et al., 2025) using a network model trained with DeepLabCut (Lauer et al., 2022; Mathis et al., 2018). Five body parts of each animal were tracked across all frames of the videos: nose, both ears, neck, and base of the tail. Position data were smoothed across frames using a 10 frame (i.e., 1/6 s) length pseudo-gaussian (fastsmooth function in Matlab).

### Behavioral events

To identify the timing of behavioural events, including social interactions and object sniffing, BORIS (Behavioral Observation Research Interactive Software;(Friard C Gamba, 2016) was used to quantify the amount of time spent performing each behaviour and when it was taking place. Our ethogram included sniffing of another animal’s nose or mouth (“face-to-face”), anogenital investigation (“rear-sniff”), sniffing of the body or tail (“body sniffing”), and a set of competitive or play behaviors such as mounting, wrestling, biting, boxing, chasing, and nape attack (“agonistic”). Scoring was performed by 4 individuals blind to lesion group who received videos in a pseudo-randomly selected way that controlled for number of videos in each experimental group. To maintain inter-rater reliability, author ML scored the majority of the videos, and then trained and continued to supervise the 3 other scorers.

### Analysis

Scored data from BORIS was analyzed using custom-written code in MATLAB (Mathworks; Nantick, MA).

Relative orientation, movement, and position (ROMP) variables were calculated as follows: Direction-to-other was computed as the 4-quadrant arctangent (Matlab arctan2 function) of the vector described by center of head (average position of neck and both ears) to nose, relative to the angle between the center of head to the stimulus animal’s nose (or tail-base in analysis of rear-sniffing). Movement calculations focused on the animal’s linear velocity, as computed by the change in nose position over time, or relative velocity as the change in distance between two individuals (typically nose-to-nose) over time. Relative position (distance) estimates similarly included both the nose position relative to the center of chamber and nose position relative to another individual’s nose (or tail-base in analysis of rear-sniffing). To address predictions, we used up to 11 measures: the slope of rats’ deceleration, their highest relative speed, the closest distance (i.e., distance at the computed onset of interaction), their overall approach speed, the average speed before approach, the partner’s speed during approach, the moment-to-moment correlation between focal and partner’s speed (i.e., coordination/synchrony of movement), the head direction (angle) of the approach, the angle described by the two partner’s body’s during approach (e.g., were approaches more often from the front, the side, the rear), the peak rotational speed during approach, and the partner’s peak rotational speed after contact was made (e.g., after onset of a rear-sniff, did the partner immediately begin turning around).

To evaluate ROMP changes in relation to a behavioral event, such as distance between two animals over a period prior to face-to-face contact, human rater events were re-aligned by using the video frame at which the body parts stop approaching and begin moving away from one-another, nearest to the BORIS-scored time for that interaction. For example, the “object-sniffing” start time would be the point where the nose ceased to approach and began moving away from the object center. This would be the lowest, absolute value of the relative velocity. (Note that we found this to be more reliable than the closest point of contact, which can happen later.)

Levels of interaction were examined using time-budget analyses, including cumulative distribution of interaction levels over session time.

Although figure illustrations show three experimental groups (RSC, PPC, sham), the experimental design did not pair RSC against PPC, so all statistical comparisons were made by separate, two-sample t-tests comparing the lesion group (n = 11) against its paired, sham-control group (n = 11). In several cases, such as comparisons for each of five behavior interaction types, it was necessary to correct for multiple comparisons. In these cases, a Benjamini-Hochberg false discovery correction was used. To avoid false-negatives when testing 11 ROMP measures during social approach, given the small sample size, an alpha of 0.01 was used and positive results were considered within the context of predictions.

Code for all analyses can be found on GitHub (https://github.com/ninsel/rsc_lesion). Video data can be made available on Databrary (https://databrary.org/), and processed data will be shared on request.

#### Transparency and openness

We report how we determined our sample size, all data exclusions, all manipulations, and all measures in the study. Code for all analyses can be found on GitHub (https://github.com/ninsel/rsc_lesion). All videos are openly available on Databrary (https://nyu.databrary.org/volume/1982), manually scored and tracked data files will be shared on request. Data were analyzed using MATLAB R2024a. Neither the study’s design not analyses were pre-registered.

## RESULTS

### Effects of lesions on movement and spatial behavior

Ibotenic acid lesions caused visible loss of tissue in the PPC in the PPC group (mean = 5.3 mm^3^ of the 11.8 mm3 region, or 45.1%) and RSC in the RSC group (mean = 11.4 mm^3^ of the 18.6 mm^3^ target region, or 61.1%; **Figure 2**). Although this suggests only partial lesions to the regions, the tissue loss likely underestimates the extent of damage, as exposure to the excitotoxin can cause inflammatory processes that damage neurons beyond the site of tissue loss (Marty et al., 1991; Roffman, 2000). Even use of NeuN markers over the presently used Nissl stain has been shown to reveal lesions nearly twice the volume (Cholvin et al., 2018).

**Figure 2.**
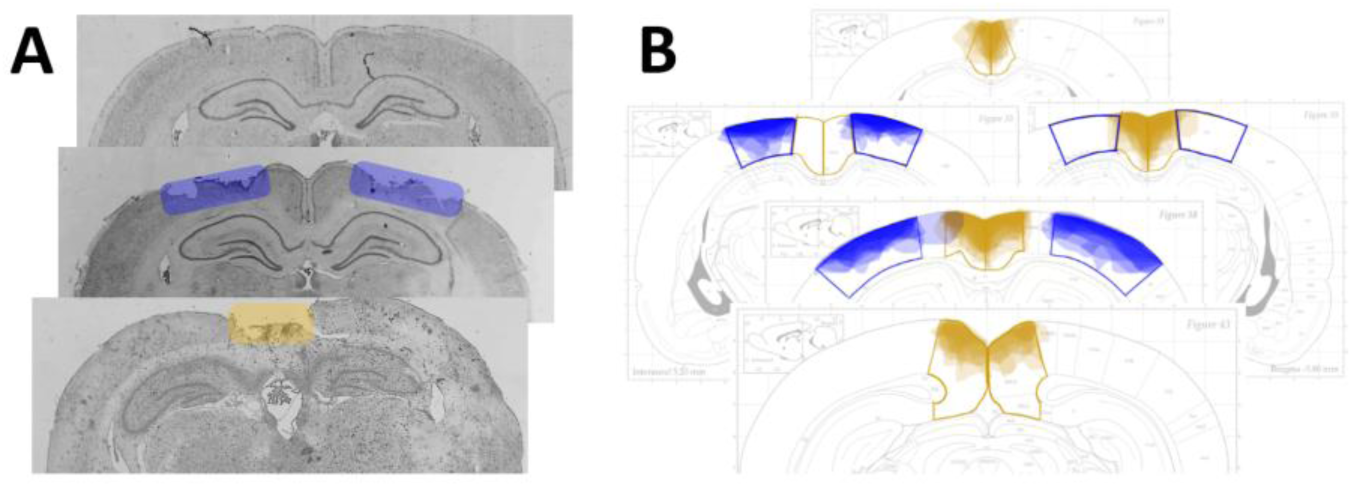
Lesions extent. *Note*. (A) Representative histological sections from sham (top) PPC (middle) and RSC (bottom) rats. Blue shading shows PPC target region in this coronal plane—in this case superficial tissue was lost and deeper layers showed some damage and some sparing. Orange shading shows RSC target in this plane; nearly all tissue was ablated by ibotenic acid. (B) Each panel shows a coronal section from the rat brain atlas taken from different coronal planes along the anterior-posterior axis. Box outlines illustrate the full target region (blue, parietal; orange, retrosplenial) and shading indicates the extent of visible tissue loss from each animal (darker shaded indicates overlap across more individuals).

Lesions appeared to impact movement in the experimental chamber. Most notably, median velocity was higher in RSC lesioned animals across the five behavioral conditions (open field, social, static social, object, and dynamic object; mixed ANOVA of RSC versus paired shams, effect of lesion: F(1,20) = 6.6, p = 0.018, behavior condition: F(4, 80) = 13.1, p = 2.9 x 10^-8^ , lesion x behavior: F(4, 80) = 0.86, p = 0.49; **Figure 3A**; medians were used rather than means because they were less sensitive to bouts of speed associated with specific interactions with social or non-social stimuli RSC animals also showed a trend for faster accelerations and decelerations (lesion: F(1,20) = 3.67, p = 0.070, behavior condition: F(4,80) = 14.7, p = 4.8 x 10^-9^, lesion x condition: F(4,80) = 1.2, p = 0.31). Unlike RSC lesioned animals, PPC lesions did not result in greater movement (lesion: F(1,20) = 2.68, p = 0.12, behavior condition: F(4,80) = 12.0, p – 1.1 x 10^-7^; lesion x condition: F(4,80) = 0.97, p = 0.43); however, the direction of change was the same, with statistical trends in both social (p = 0.045; false discovery correction across conditions) and object (p = 0.057) conditions, suggesting potentially inadequate statistical power and insufficient PPC lesion spread.

**Figure 3.**
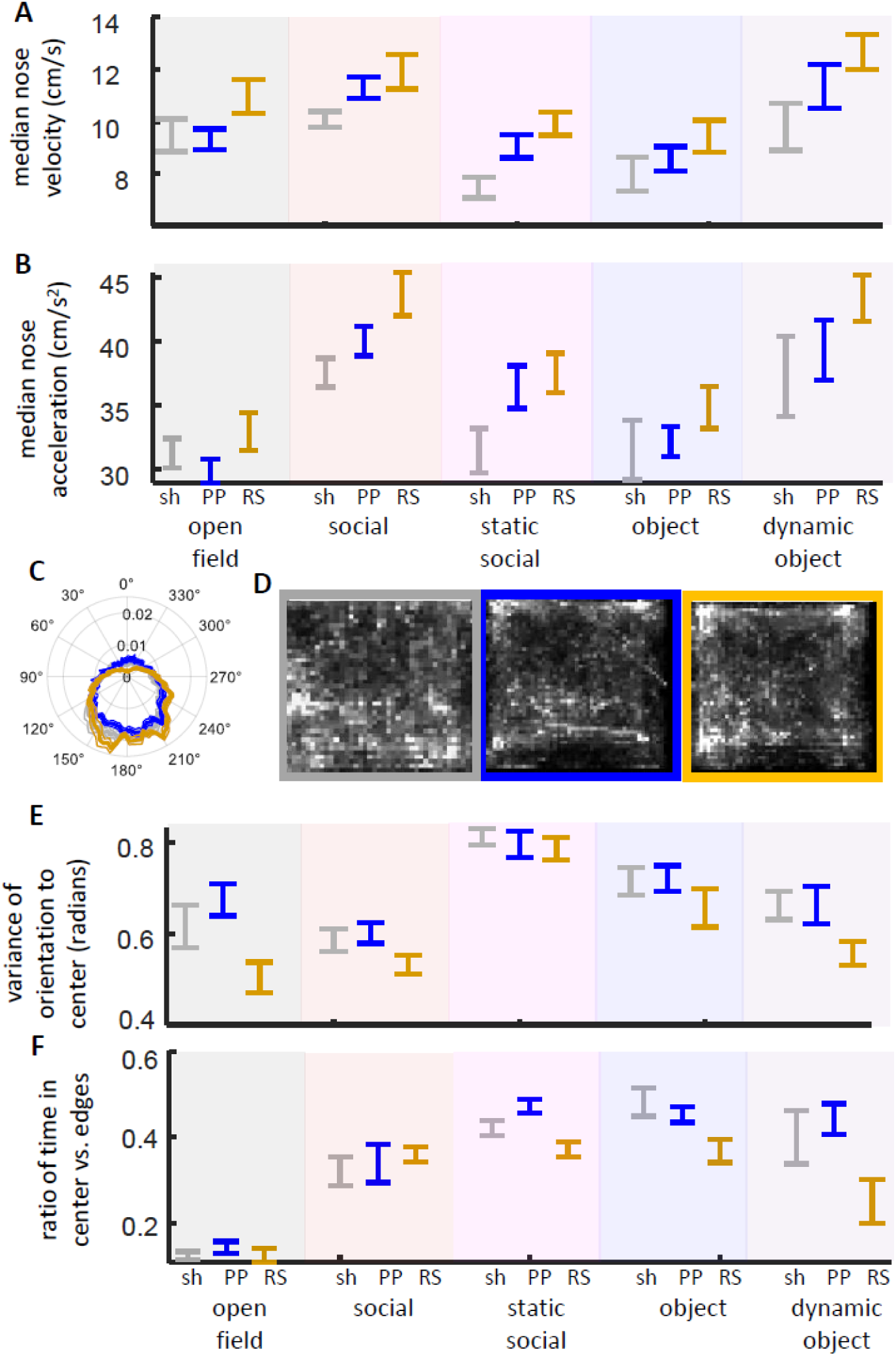
Movement and spatial behavior. *Note*. (A) Median velocity within animals during each behavioral condition, averaged within groups (±SEM). RSC animals consistently moved more during the 10 min sessions than did their paired shams. (B) Absolute value of nose movement acceleration also suggest faster movements among RSC animals. (C) Averaged distribution of head orientation toward (0°) versus away from (180°) the center of the chamber during the open field sessions (thick line = mean across rats; thin lines = SEM). RSC lesions (orange trace) reduced orienting to the center, with proportionately higher orienting between 120° and 240° away. (D) Nose position occupancy maps, averaged across animals within lesion groups, during the open field sessions (left to right: sham, PPC, RSC). (E) Circular variance for distribution of orienting toward the center, for each behavioral condition, averaged across animals within experimental groups. Lower numbers indicate less variance and therefore a stronger preference away from center. (F) Ratio of time of nose in the central versus outer half of the occupancy map.

Lesions also affected how rats oriented in space. Across behavioral conditions, RSC rats were less likely to orient toward the center of the chamber relative to the sides; i.e., circular variance of gaze direction was reduced, more concentrated away from the center (lesion: F(1,20) = 5.4, p = 0.030, behavioral conditions: F(4,80) = 39.2, p = 3.6 x 10-18, lesion x behavior: F(4,80) = 0.77, p = 0.55; **Figures 3CsE**). This was also partly reflected by a trend for different Rayleigh z-scores testing for non-uniform distributions (F(1,20) = 4.06, p = 0.057). This effect was not observed in PPC rats (p >> 0.05). There was no evidence of differences in occupancy patterns (e.g., occupancy in central vs. edge pixels; both PPC vs sham and RSC vs sham: p > 0.10; **Figure 3DsF**).

### Effects of lesions on social attention and interaction

Across all groups, rats tended to orient away from a freely-moving social stimulus more than toward (**Figure 4A**). A statistical trend suggested that this tendency to orient away from a social stimulus was stronger in RSC rats (two sample t-test of circular variance between RSC and paired sham animals: t(20) = 2.07, p = 0.051; t-test comparing Rayleigh z-scores: t(20) = 2.05, p = 0.054; **Figure 4A**). However, orienting away from a social stimulus was highly correlated with RSC rats’ tendency to orient away from the center of the experimental chamber (Pearson correlation of circular variances in RSC group: rho = 0.70, p = 0.016; **Figure 4B**). Since “away from center” orienting was observed across behavioral conditions, including the open field, the behavior was unlikely to be due to the social stimulus itself. Interestingly, there was also a correlation between social orienting of RSC and that of their stimulus partner (rho = 0.63, p = 0.034). This correlation could be found across all animals (n = 37, rho = 0.60, p = 9.0 x 10^-5^; **Figure 4C**), suggesting that social attention between animals was largely reciprocal over the course of a session.

**Figure 4.**
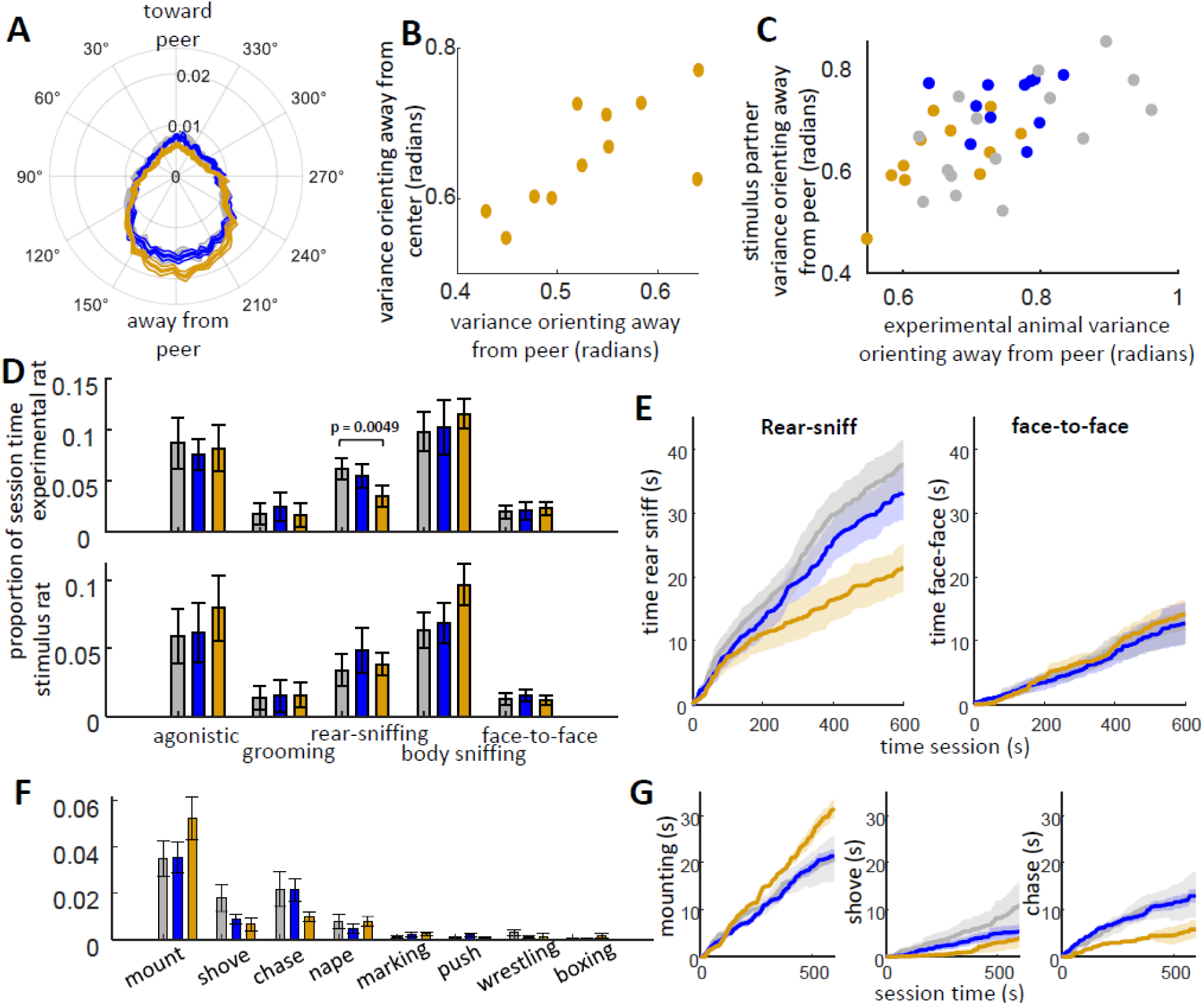
Social orienting and interaction. *Note*. (A) Circular histogram showing proportion of time experimental rats looked toward their stimulus partner (toward 0°) versus away (toward 180°). Thick lines are averages across rats, thin lines ±SEM. (B) Scatterplot showing correlation between degree to which RSC rats oriented away from center and oriented away from the stimulus partner (peer). (C) Scatterplot showing relationship between one dyad member’s degree of peer orienting and another’s. (D) Levels of interaction across 5 interaction types, as measured by proportion of session time. Top plot shows experimental (lesion) animal, bottom shows stimulus partner. (E) Cumulative distributions showing progressive amount of time that animals spent rear-sniffing (left plot) or in face-to-face (right plot). Rear-sniffing differences only began to emerge after the first half of the session. (F) Relative levels of multiple agonistic behaviors across groups. No significant differences were observed. (G) Follow-up exploratory analyses on cumulative differences between mounting, shoving, and chasing over session time.

No compelling differences were observed in the degree or type of social interaction lesion groups engaged in (**Figure 4B**). Across 5 behavioral categories, RSC rats only showed a reduction in rear-sniffing (t(20)= 3.16, p = 0.0049, significant with Benjamini & Hochberg false discovery correction across 5 tests). Further analysis revealed these differences only began to emerge after the first 5 minutes, suggesting both the ability and initial interest in anogenital sniffing were intact (**Figure 4E**). A separate set of analyses compared specific competitive (agonistic/play) behaviors, including mounting, shoving, chasing, nape attacks, marking, rear-pushes, wrestling, and boxing (**Figure 4F**). No statistical differences were observed (p > 0.1), though the use of these behaviors across rats was variable. Further exploratory investigation was made into shoving and chasing based on potential decreases in RSC rats (p < 0.2). While there were hints that shoving was reduced among RSC rats during early stages of the social period (e.g., p < 0.05 at 5 minutes; **Figure 4G**), the observation was unrelated to hypotheses and did not pass statistical criteria. Additional analyses examining effects of lesions on non-social behaviors (self-grooming and investigative rearing) found no differences between groups.

### Effects of lesions on social approach and movement

The primary prediction of the present study was that lesions to the RSC or PPC would impact how rats approached one-another during freely moving social interaction. We examined how animals’ relative orientation, movement, and position (ROMP variables) changed in relation to the onset of interaction events, with a focus on face-to-face and rear-sniffing (chosen because both interaction types are well classified, but likely have different meaning and involve distinct dyadic poses). Many measures focused on the time between when an animal began approaching its partner (either the final point at which relative distance began to decrease or, if there was no final point of decrease within the 5 seconds, then the point within 3 s at which the rat began decelerating toward its partner) and when interaction “contact” was made (as defined by when the distance between relevant body parts—e.g., nose and nose for face-to-face, or nose and tail-attachment for rear-sniffing—were no longer approaching). This time period averaged about 0.83 s (0.63 standard deviation) for face-to-face and 0.65 s (0.56 standard deviation) for rear-sniffing. **Table 1** lists all ROMP measures taken around the time of interaction events, along with statistical comparisons (two sample t-tests, alpha = 0.01—see METHOD) between RSC and sham groups after taking within-animal averages. Average ROMP traces around interaction time are presented in **Figure 5**. No systematic differences were observed between PPC and paired shams. RSC rats similarly showed almost no differences relative to shams, with a few minor exceptions. There were statistical trends (0.05 > p > 0.01) for faster linear and rotational approach speed before rear-sniffing—consistent with prior observations for overall greater movement among RSC animals. The strongest difference observed was not in the experimental animals, but in the stimulus partners: when stimulus partners performed a rear-sniff, they moved much more quickly with RSC compared with the paired, sham controls. This is also consistent with overall greater movement among RSC animals: i.e., stimulus animal rear-sniffs were more likely to take place as the lesioned rat was moving. This was confirmed by a follow-up analysis: a higher ratio of rear-sniffs performed by the stimulus animal took place when their partner was moving faster than 7.5 cm/s in the RSC group compared with paired shams (t(20) = 2.8, p = 0.010; also true for the PPC group: t(20) = 2.12, p = 0.047). To summarize: we found no evidence that socially-directed movements were impacted by the lesions, only that overall changes in movement (increased velocity) in RSC rats altered the dynamics of stimulus partner rear-sniffs.

**Figure 5.**
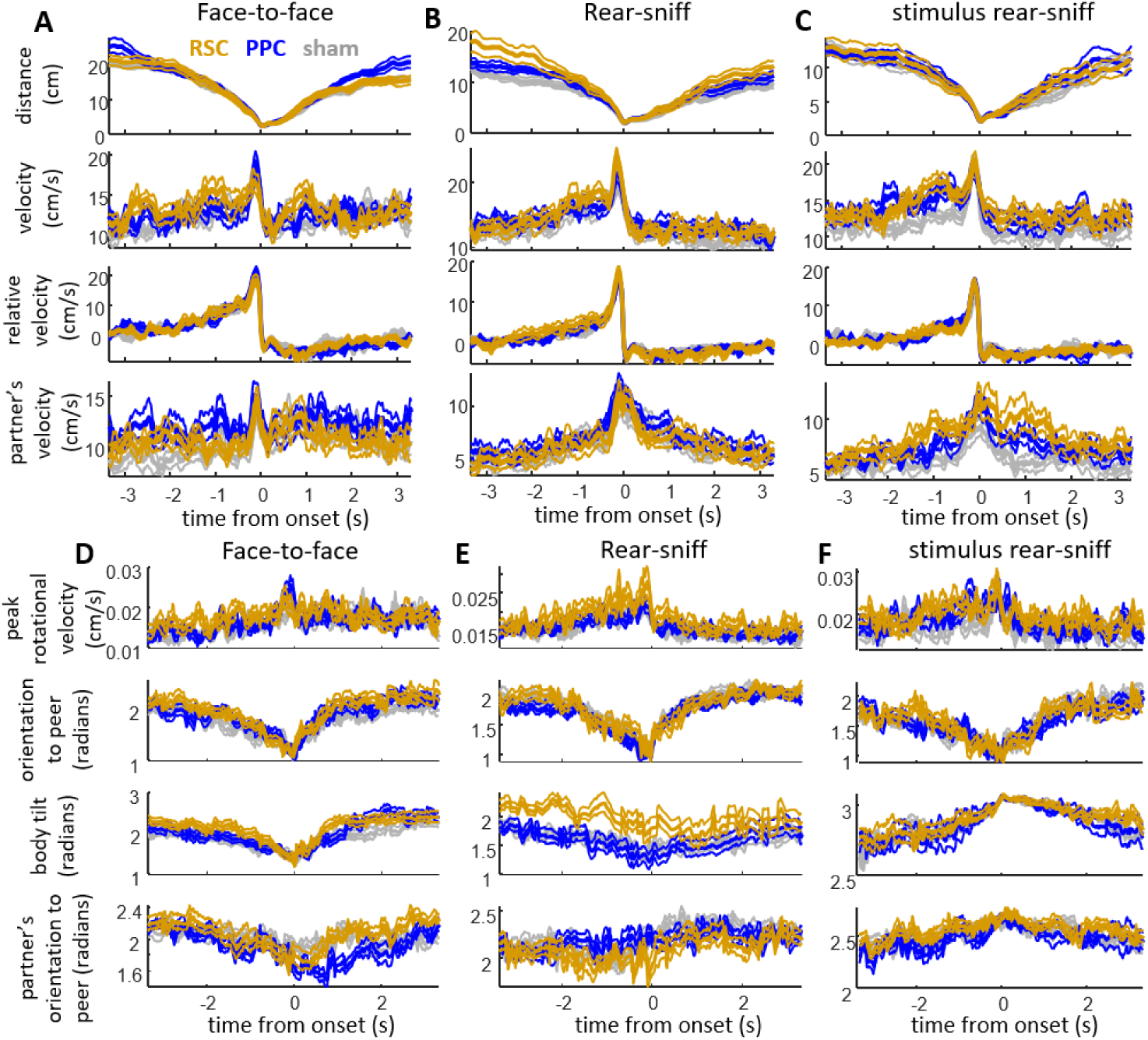
ROMP measures before and after social interactions. *Note*. A) Linear distance and movement variables aligned to the onset of face-to-face interactions, averaged within sessions and then across animals for each group. Variables from top to bottom: distance between each nose, speed of animal, relative speed, and speed of the stimulus partner; RSC orange, PPC blue, shams gray). Traces show levels of each variable over time relative to the onset of interaction. (B) As in part A, but aligned to onset of rear-sniffs. Here, distances and relative velocity are between experimental animal’s nose and stimulus-animal’s tail-attachment. (C) As in part B, but aligned to the onset of stimulus-animal’s rear-sniffs. Distances and relative velocity are between stimulus animal’s nose and experimental animal’s tail-attachment. (D-F) Averaged, angular measures and rotational movement aligned to the onset of face-to-face (D), rear-sniffing (E), and partner’s rear-sniff (F) interactions. From top to bottom: peak rotational velocity (i.e., change in head direction), head orientation to peer (0 is directed toward stimulus animal, π is directed fully away), angle between each body (0 is animal bodies facing toward one-another, π is facing in the same direction) and partner’s head-direction (0 toward, π away).

**Table 1.** Statistical comparisons between RSC and paired shams across ROMP measures.

|  | Face-to-face | Rear-sniff | Other's rear-sniff |
| --- | --- | --- | --- |
| Slope of approach deceleration | $t = 1.79$<br>$p = 0.65$ | $t = 1.46$<br>$p = 0.16$ | $t = 0.92$<br>$p = 0.37$ |
| Highest relative speed | $t = 1.03$<br>$p = 0.31$ | $t = 0.86$<br>$p = 0.40$ | $t = 1.15$<br>$p = 0.26$ |
| Closest distance reached | $t = 0.75$<br>$p = 0.46$ | $t = 1.06$<br>$p = 0.30$ | $t = 0.44$<br>$p = 0.66$ |
| Approach speed | $t = 1.28$<br>$p = 0.22$ | $t = 2.20$<br>$p = 0.040$ | $t = 2.96$<br>$p = 0.0077$ |
| Partner's speed during approach | $t = 0.51$<br>$p = 0.62$ | $t = 0.15$<br>$p = 0.88$ | $t = 1.57$<br>$p = 0.13$ |
| Velocity correlation with partner (synchrony) | $t = 0.35$<br>$p = 0.73$ | $t = 1.14$<br>$p = 0.27$ | $t = 0.54$<br>$p = 0.60$ |
| Speed before approach | t = 1.26<br>p = 0.22 | t = 0.77<br>p = 0.45 | t = 3.00<br>p = 0.0071 |
| Directionality of approach | t = 1.21<br>p = 0.24 | t = 1.16<br>p = 0.26 | t = 0.72<br>p = 0.48 |
| Angle between two partner's bodies | t = 0.83<br>p = 0.41 | t = 1.85<br>p = 0.079 | t = 2.04<br>p = 0.055 |
| Peak rotational speed during approach | t = 0.49<br>p = 0.63 | t = 2.55<br>p = 0.019 | t = 0.60<br>p = 0.56 |
| Peak rotational speed after contact | t = 0.12<br>p = 0.91 | t = 0.99<br>p = 0.33 | t = 0.17<br>p = 0.86 |

### Effects of lesions on object, static social, and dynamic object interactions

The negative results observed in ROMP measures during social interaction were reflected by similarly negative results in the conditions with control stimuli. In the object test, groups did not differ in the degree of sniffing (PPC: t(20) = 1.46, p = 0.16; RSC: t(20) = 0.39, p = 0.70; **Figure 6AsB**); nor in any ROMP measure of object approach (velocity of approach, duration of approach, closest distance to object after initial approach, average orienting during approach: p >> 0.10; **Figure 6C**). The same was true in the static-social condition, both in the degree of sniffing (PPC: t(20) = 1.07, p = 0.30; RSC: t(20) = 0.26, p = 0.79; **Figure 6DsE**) and ROMP measures of approach (velocity of approach, duration of approach, closest distance to stimulus after approach, average orienting during approach: p >> 0.10 except for closest distance in PPC, p = 0.069, non-significant and unrelated to other conditions or hypotheses; **Figure 6F**). In follow-up exploratory analyses, we found hints of reduced object investigation among PPC rats during early parts of the session (p < 0.05 between 84 and 132 s; **Figure 6B**), although there were no differences in median inter-sniff intervals (Wilcoxon signed-rank test, p >> 0.1 for both PPC and RSC), and clearly no differences between groups in investigation of the static social stimulus (**Figure 6F**).

**Figure 6.**
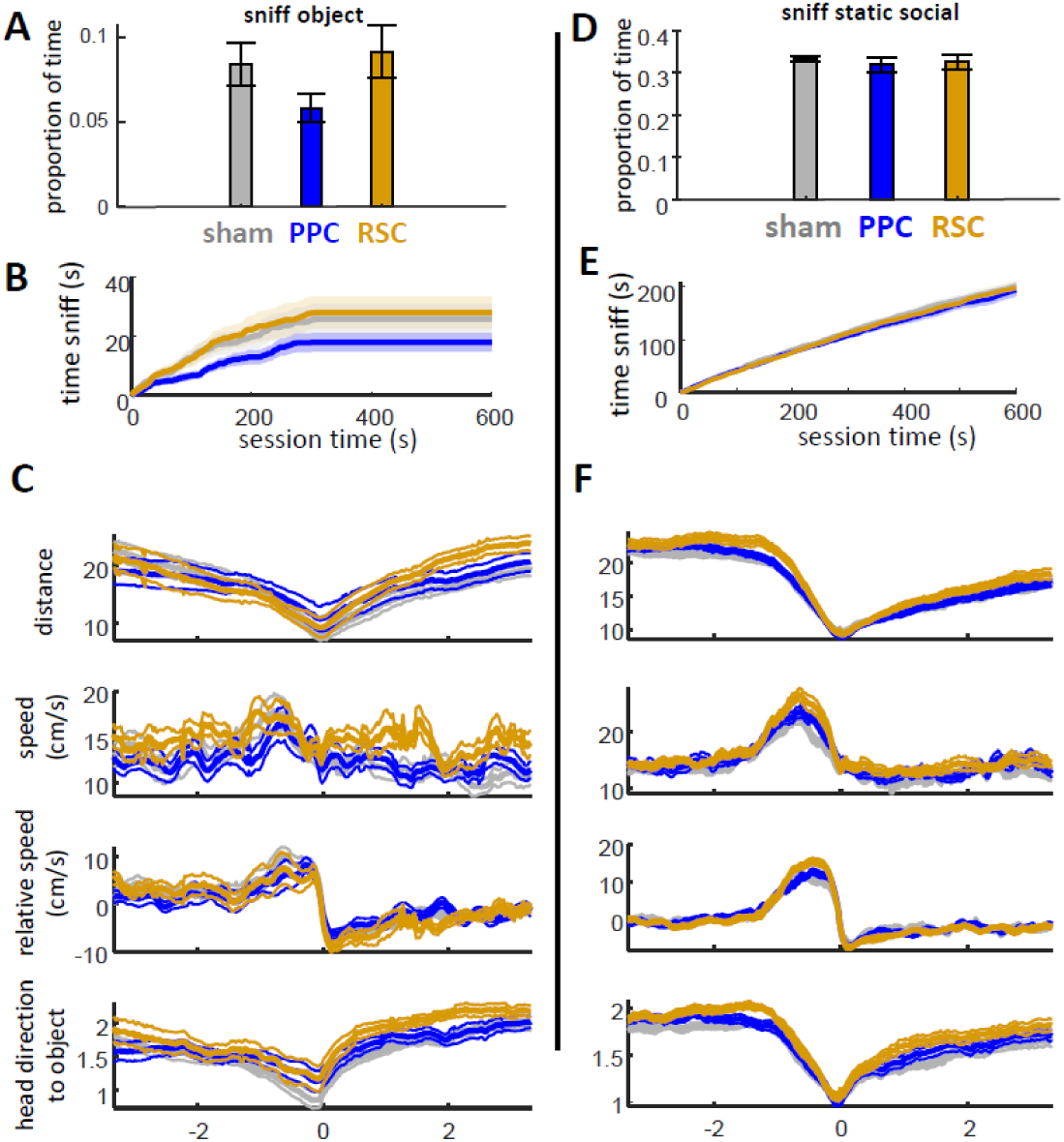
Behavior during object and static-social conditions. *Note*. (A) Levels of sniffing across experimental animals, as measured by proportion of session time. (B) Cumulative levels of sniffing throughout the session. (C) From top to bottom, distance, speed, velocity relative to object, and head direction relative to object aligned to sniff onsets, averaged within sessions and then across animals (thicker lines are means, thinner lines ±SEM). (D) Levels of sniffing of the static social stimulus (note change in scale from A). (E) Cumulative levels of sniffing throughout the session. (F) Averaged distance, speed, relative velocity, and head-direction aligned to sniff onsets.

In the dynamic object condition, 3 different types of behaviors were examined (chase, sniff, avoid), as well as movement responses to a stimulus behavior (“bump”—when the moving stimulus makes contact against the rear or body of the rat). Unexpectedly, chases appeared to be higher in the PPC group (chase, PPC: t(20) = 3.94, p = 8.1 x 10^-4^, RSC: t(20) = 1.23, p = 0.23; sniffs, PPC: t(20) = 1.23, p = 0.23, RSC: t(20) = 0.58, p = 0.57; **Figure 7A**). There was also a statistical trend for PPC rats to show reduced avoidance (PPC: t(20) = 2.02, p = 0.057; RSC: t(20) = 0.63, p = 0.54). This appeared to be due to longer chase durations (PPC: t(20) = 3.1, p = 0.0055; RSC: t(20) = 1.05, p = 0.31). They also showed a statistical trend for longer sniff durations (PPC: t(20) = 1.92, p = 0.070). However, PPC rats did not appear to orient more toward the moving stimulus (comparison of distribution variances, PPC: t(20) = 0.18, p = 0.86, RSC: t(20) = 1.24, p = 0.23; **Figure 7B**). When examined over session time, the differences between lesion groups appeared to emerge during the second half of the session (**Figures 7C-E**). RSC rats, in contrast, only showed statistical trends for faster peak velocities during chase (PPC: t(20) = 0.51, p = 0.55; RSC: t(20) = 2.01, p = 0.058; **Figure 7F**) and faster velocities following a bump by the moving object (PPC: t(20) = 0.45, p = 0.66, RSC: t(20) = 1.82, p = 0.084; **Figure 7H**)—consistent with prior results suggesting overall increased movement in this group. Otherwise, no systematic differences were observed between groups with respect to interaction-related ROMP variables (including closest touch, rat orienting toward the stimulus, or movement speeds of the human-controlled moving stimulus, p >> 0.10).

**Figure 7.**
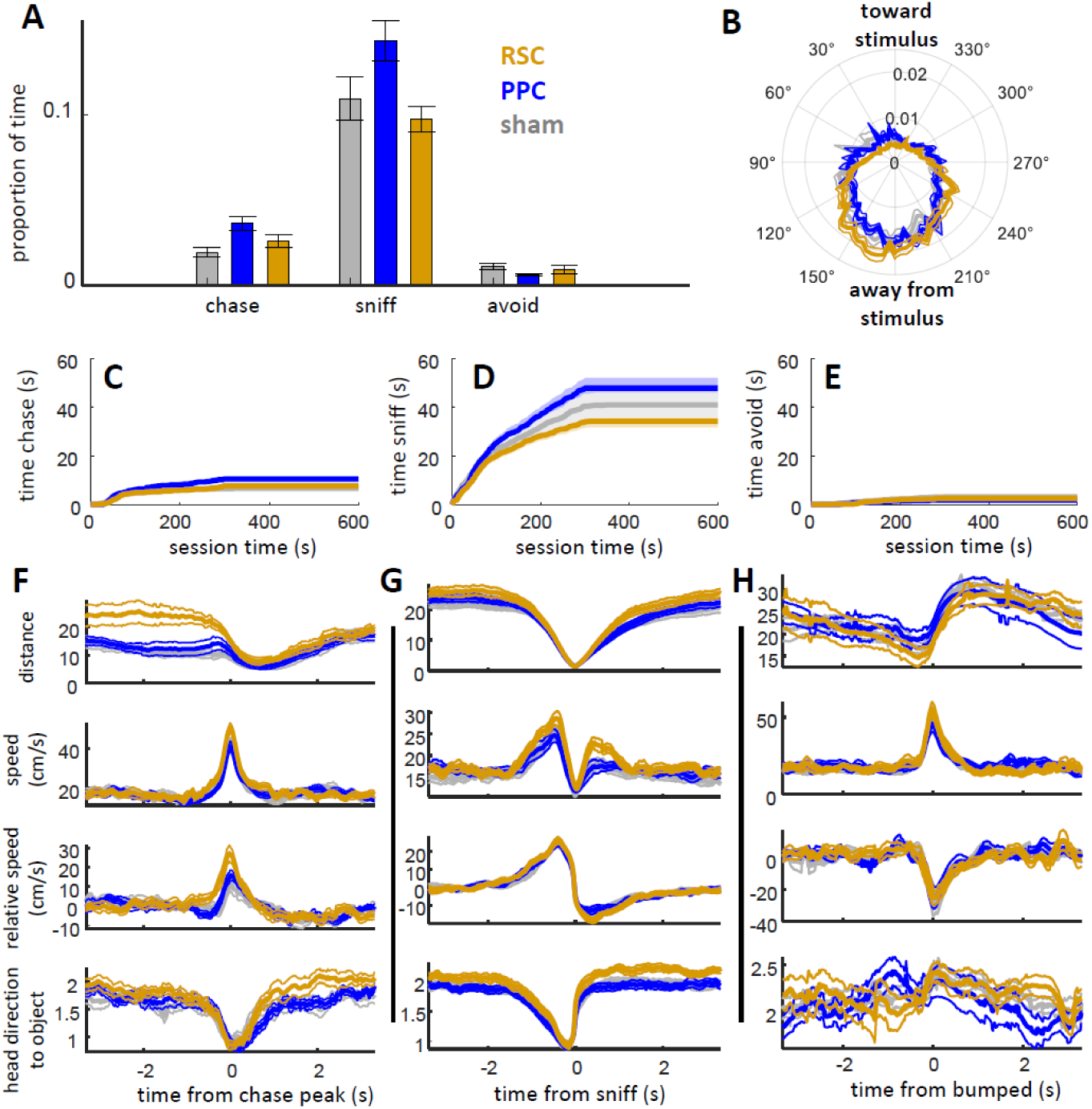
Behavior during dynamic object condition. *Note*. (A) Levels of chase, sniff, and avoid of the dynamic object. (B) Relative degree of orienting toward (near 0°) versus away from (near 180°) the dynamic object. (C-E) Cumulative time chasing (C), sniffing (D), and avoiding (E) the dynamic object. (F) Distance, speed, velocity relative to the object, and head relative direction to the dynamic object aligned to the peak velocity of chase events (averaged across events, across rats; thin lines are ±SEM across rats). (G) As in part F, aligned to the onset of sniffs. (H) as in part F, aligned to when objects bumped against the experimental animal.

## DISCUSSION

Here we tested the hypothesis that the PPC and RSC are involved in processing and managing distance with another individual, and that lesions to these regions in rats would therefore alter social approach patterns. Although lesions to the RSC caused generalized increases in movement, along with more time orienting away from the center of the chamber, we found no evidence that the dynamics of social approach changed relative to non-lesioned (sham) controls. Lesions to the PPC appeared to increase the degree to which these animals chased a moving object stimulus, including increasing the duration of the chase, but otherwise showed no consistent differences relative to shams. While the mild effects of PPC lesions could be due to tissue loss in only part of the region (particularly superficial layers), the data generally suggest a limited role of these regions in the sensorimotor dynamics associated with spontaneous social approach—at least within the open, 50 cm^2^ space of the experimental chamber. It may therefore be that the RSC and PPC are differentially required when social interaction involves greater cognitive demands, including larger, more complex environments, or environments with multiple dynamic stimuli, in conditions that depend more strongly on predictions, attention, and the use of multiple spatial frameworks.

Lesions did not appear to have selective effects on social-approach patterns, but more general effects on behavior had some indirect effects on social engagement. Put another way: all changes related to the dynamically moving stimulus (e.g., a peer) could be better described by changes in how rats treated static features of the environment. For example, RSC rats tended to orient way from their stimulus partner, which at first glance could be taken to mean that RSC rats were avoiding their peer. This was, however, highly correlated with rats’ tendency to look away from the chamber center more generally, which we observed even in non-social conditions. Similarly, lesioned rats were more likely to be running faster when receiving a rear-sniff from their stimulus partner—almost as though escaping the investigation. But lesions, particularly to the RSC, also caused increase movement in non-social settings, and it seems more likely that the “social” result reflected the stimulus animal having to catch-up when investigating from behind.

This lack of evidence for selectively social changes offers a different perspective from prior research findings, which suggest that changes of the RSC underly some, apparently “social” autism-like symptoms in certain genetic mouse lines (Garrido et al., 2022; Shang et al., 2021; Yang et al., 2021). Notably, these prior studies do not implicate the RSC in social interaction per se, but point to a role of the region in relative investigation of novel compared with familiar social stimuli. Since social context effects were not the focus of the present investigation, rats were not offered a choice between novel and familiar peers; however, RSC lesions were associated with reduced rear-sniffing during the latter 5 minutes of the social interaction sessions, relative to sham controls, suggesting that salience to the novel peer diminished more rapidly. This temporal pattern is similar to the changes in interaction levels in another rodent species when presented with familiar compared with novel social stimuli, where relative decreased interaction between familiar dyads emerge only after the first 5 minutes (Figure 4D of Lidhar et al., 2021). The dynamics of RSC involvement in social investigation may also be more subtle. A recent report suggests the mouse RSC exhibits an “activation-to-inhibition” pattern upon social contact that, when manipulated experimentally, alters social investigation (Shi et al., 2024). It is difficult to know the extent to which these phenomena should be considered in a strictly social context, relative to stimulus salience or the processing of self-motion and space.

The absence of changes in social approach, as measured by relative orientation, movement, and position (‘ROMP’) variables, was unexpected. In humans and other primates, the PPC has been shown to be important for animals to adjust movement according to unexpected sensory changes, based on behavioral changes following damage to the region (Gréa, 2002; Pause et al., 1989; Pisella et al., 2000) and as inferred from physiological patterns (Buneo & Andersen, 2006; Freedman & Ibos, 2018). Physiological studies have also pointed to a role of the PPC not in online-adjustments per se, but in maintaining expectation of one’s own future position/movement (Li et al., 2022). These studies focus on hand movements and reach, rather than whole-body social proxemics. Given the apparent role of the PPC in navigational movements (Alexander et al., 2022; Nitz, 2006; Whitlock, 2017; Whitlock et al., 2012; Wilber et al., 2014), it seems likely that reach-related functions studied in primates extend to whole-body movement planning and updating. One potential reason we failed to detect social approach differences is that the lesions did not sufficiently extend into areas of the PPC involved in inter-individual space; however, as noted in Results, loss of tissue likely underestimates the spread of neural damage (Cholvin et al., 2018). In fact, rather than disrupting approach behaviors, PPC animals increased the degree to which they chased the dynamic, non-social object, and there was a statistical trend for reduced object avoidance. These patterns were not found in the social conditions, perhaps highlighting meaningful differences between ball-chasing and social approach. All together, it seems unlikely that the dorsal regions of PPC and RSC that were lesioned in the present study are required for simple sensorimotor adjustments during nearby social approach. Social approach may be supported by multiple redundant networks involving brainstem and cerebellum, with the PPC adjusting these based on expectations and maintained plans (as discussed by Li et al., 2022). PPC and RSC may be only be necessary when more complex integration of information is required, such as when animals may be maintaining information about a potential target (as suggested by an RSC role in guiding animals to an escape; Campagner et al., 2023) or when making navigational approach decisions according to peer salience (e.g., Dautan et al., 2024; Garrido et al., 2022; Shang et al., 2021; Shi et al., 2024; Yang et al., 2021).

It therefore remains unclear how functions of the RSC change when spatial targets dynamically change their spatial locations. The present study’s use of dynamic stimuli has to be considered within the restricted spatial environment of a 50 cm^2^ open field, and the absence of social decisions beyond to a single, unfamiliar, same-sex individual. It also may be constrained by the scope of the lesions: for both RSC and PPC, lesions were made to more dorsal areas. In primates, perception of social-movement, and the inferences that stem from social movements, are processed along more lateral areas of cortex (Pitcher C Ungerleider, 2021); and, in rodents, responses to moving stimuli have been found in more temporal “ectorhinal” cortex (Nishio et al., 2018). Having now established potentially useful analytic approaches, the research can be further developed to include complex environments in which animals have to maintain expectations of others’ probable locations without clear lines of site; to multi-animal conditions in which one individual is more socially salient than another, and to neural manipulations that target a broader (including more ventral) scope of the cortex. The value of the present study may therefore be largely in developing a pipeline for multi-animal tracking and ROMP measures, which can be further applied to more elaborate examinations of natural behaviour.

Our use of ROMP measures builds on a history of investigations on inter-individual distances in animals (Hediger, 1955) and humans—classically referred to as “proxemics” (Hall, 1966)—along with more contemporary, neuropsychiatric considerations of interpersonal space (Lee et al., 2024). Neural descriptions of interpersonal space (“neuroproxemics”) often suggest a role of the “frontal-parietal” network but, perhaps owing to spatial constraints of human neuroimaging methods, typically focus on intrusions of personal space (Basile et al., 2024; Cartaud et al., 2018), but see also (Bufacchi & Iannetti, 2018). Understanding how individuals manage inter-personal space will also require more than analysis of inter-individual distance (*relative position*)—it will have to include how individuals direct attention to one another (typically measurable through *relative orientation*) and the dynamics of their *relative movements* across distances and time from a social interaction. It has become increasingly possible in animal research to capture these metrics thanks to advancements in computer vision, along with associated software packages enabling tracking of multiple body parts (pose estimation) in more than one animal at a time (Chen et al., 2020; Coulombe et al., 2024; Lauer et al., 2022; Pereira et al., 2022; Romero-Ferrero et al., 2019; Walter & Couzin, 2021). These tools still often mis-identify similar-looking, closely-interacting individuals, so we developed and used DIPLOMAT (Robinson et al., 2025) to help reduce body-swap errors and improve manual editing. The present work helps establish methods that can be leveraged for future neuroproxemics research.

The results speak against high-level parietal/retrosplenial regions being *necessary* for sensorimotor behaviors associated with social approach. This does not mean the regions do not process this information, or that they don’t use it, when available, during online approach or more complex social-spatial behaviors. These negative results therefore offer one step toward better understanding the ways in which these traditionally “spatial” regions are recruited in social contexts.

## ACKNOWLEDGEMENTS

Research was supported by NSERC grant RGPIN-2023-05624. We thank Rhiannon Izatt and Letitia Collins for assistance with data collection, Rhiannon Izatt and Menahil Ayub for assistance with data processing, and Isaac Robinson for frequent updates to DIPLOMAT.

